# Horizontal transfer and subsequent explosive expansion of a DNA transposon in sea kraits (*Laticauda*)

**DOI:** 10.1101/2021.06.13.448261

**Authors:** James D. Galbraith, Alastair J. Ludington, Kate L. Sanders, Alexander Suh, David L. Adelson

**Affiliations:** School of Biological Sciences, University of Adelaide, Adelaide, SA 5005, Australia; School of Biological Sciences, University of East Anglia, Norwich Research Park, NR4 7TU, Norwich, United Kingdom; Department of Organismal Biology - Systematic Biology, Evolutionary Biology Centre, Uppsala University, SE-752 36 Uppsala, Sweden

**Keywords:** horizontal transfer, transposable element, Serpentes

## Abstract

Transposable elements (TEs) are self replicating genetic sequences and are often described as important “drivers of evolution”. This driving force is because TEs promote genomic novelty by enabling rearrangement, and through exaptation as coding and regulatory elements. However, most TE insertions will be neutral or harmful, therefore host genomes have evolved machinery to supress TE expansion. Through horizontal transposon transfer (HTT) TEs can colonise new genomes, and since new hosts may not be able to shut them down, these TEs may proliferate rapidly. Here we describe HTT of the *Harbinger-Snek* DNA transposon into sea kraits (*Laticauda*), and its subsequent explosive expansion within *Laticauda* genomes. This HTT occurred following the divergence of *Laticauda* from terrestrial Australian elapids ~15-25 Mya. This has resulted in numerous insertions into introns and regulatory regions, with some insertions into exons which appear to have altered UTRs or added sequence to coding exons. *Harbinger-Snek* has rapidly expanded to make up 8-12% of *Laticauda* spp. genomes; this is the fastest known expansion of TEs in amniotes following HTT. Genomic changes caused by this rapid expansion may have contributed to adaptation to the amphibious-marine habitat.

## Introduction

Transposable elements (TE) are selfish genetic elements that mobilize themselves across the genome. A substantial proportion of eukaryotic genomes is composed of TEs, with most reptilian and mammalian genomes comprising between 30 and 60%. As TEs proliferate within a genome, most insertions will be either neutral or deleterious [1]. However, over evolutionary timescales the movement of TEs can enable major adaptive change; being exapted as coding and regulatory sequences, and by promoting both inter- and intra-chromosomal rearrangements such as segmental duplications, inversions and deletions through non-allelic homologous recombination [2,3].

TE expansion can also be harmful, driving eukaryotes to evolve various defence and regulatory mechanisms. Genomic shocks can disrupt this regulation, allowing TEs to expand [4]. One example of a shock is horizontal transposon transfer (HTT), in which a TE jumps from one species to another. While the exact mechanisms of HTT are unknown, many instances across eukaryotes have been reported [5–9]. Following HTT the expansion of new TEs is quickly slowed or halted due to the potentially deleterious effects they can cause [1,10], and any continued expansion will likely be slow. For example, following ancient HTT events the BovB retrotransposon has taken 32-39 My and 79-94 My for these elements to colonise between 6 and 18% of ruminant and Afrotheria genomes, respectively [6,11,12]. However rapid expansion of TEs following HT has previously been noted in *Myotis* bats, where *hAT* transposons expanded to cover 3.3% of the genome over the space of 15 Mya [13–15].

Here we report the HT of a *Harbinger* DNA transposon, *Harbinger-Snek*, into *Laticauda*, a genus of marine snakes which diverged from terrestrial Australian snakes 15-25 Mya [16–18]. Surprisingly, none of the available terrestrial animal genomes contained any trace of Harbinger-Snek, with highly similar sequences instead identified in sea urchins. Since diverging from terrestrial snakes *Laticauda* transitioned to amphibious-marine habits, foraging on coral reefs and returning to land only to digest prey, mate and lay eggs [19]. Due to the absence of *Harbinger-Snek*-like sequences from terrestrial species and highly similar sequences present in marine species, we propose *Harbinger-Snek* was horizontally transferred to *Laticauda* from a marine donor genome by habitat transition. Furthermore, since this initial HTT event, *Harbinger-Snek* has expanded rapidly within the genomes of *Laticauda* and now accounts for 8% of the *L. laticaudata* assembly and 12% of the *L. colubrina* assembly.

## Methods

All scripts/code used at: https://github.com/jamesdgalbraith/Laticauda_HT

### *Ab initio* repeat annotation of elapids

Using RepeatModeler2 [20] we performed *ab initio* annotation of the four Austro-Melanisian elapid genomes: *Laticauda colubrina* [21], *Notechis scutatus, Pseudonaja textilis,* and *Aipysurus laevis* [22]. To improve the RepeatModeler2 libraries we manually classified consensus sequences over 200 bp using a BLAST, extend, align and trim method, described by Galbraith et al. [23].

### Identification of horizontal transfer and potential source/vectors

To identify any TEs restricted to a single lineage of elapid, we searched for all TEs identified by RepeatModeler2 using BLASTN (-task dc-megablast) [24] in the three other assemblies, as well assemblies of the Asian elapids *Naja naja* [25] and *Ophiophagus hannah* [26]. TEs present in high numbers in a species, but not present in the other elapids, were considered potential HTT. This yielded a high copy number of *Harbinger* elements in *L. colubrina*. To rule out contamination, we searched for this element in a *L. laticaudata* genome assembly from GenBank. Using RPSBLAST [27] and the Pfam database [28] we identified *Harbinger* copies with intact protein-coding domains. To identify potential source or vector species, we searched all metazoan RefSeq genomes with a contig N50 of at least 10 kbp with BLASTN (−penalty −5 −reward 4 −out −word_size 11-gapopen 12-gapextend 8). In species containing similar elements, we created consensus sequences using the aforementioned BLAST, extend, align and trim method. As we had identified similar *Harbinger* elements in fish, bivalves and echinoderms from RefSeq, we repeated this process for all GenBank assemblies of other species from these clades with a contig N50 of at least 10 kbp.

We identified transposase domains present in curated *Harbinger* sequences and all autonomous *Harbinger* elements available from Repbase [29] using RPSBLAST [27] and the Pfam database. Using MAFFT (--localpair) [30] we created a protein multiple sequence alignment (MSA) of identified transposase domains. After trimming the MSA with Gblocks [31] we constructed a phylogenetic tree using FastTree [32] and from this tree chose an appropriate outgroup to use with curated elements. We subsequently constructed a protein MSA of the curated transposases using MAFFT, trimmed the MSA with Gblocks and created a phylogeny using IQ-TREE 2 (−m MFP −B 1000), which selected TVMe+I+G4 as the best model [33–35]. For comparison we also created phylogenies using the same MSA with MrBayes and RAxML [36,37]. To compare the repeat and species phylogenies, we created a species tree of major sampled animal taxa using TimeTree [38].

### Potential interaction of *Harbinger-Snek* with genes

Using the improved RepeatModeler2 libraries and the Repbase (-lepidosaur) library, we used RepeatMasker [39] to annotate the two species of *Laticauda*. Using Liftoff [40] we transferred the *No. scutatus* gene annotation from RefSeq [41] to the *L. colubrina* and *L. laticaudata* genome assemblies. To identify *Harbingers* in genes, exons and regulatory regions we intersected the RepeatMasker intervals and transferred gene intervals using plyranges [42]. To test for potential effects of these insertions on biological processes and molecular functions in *Laticauda* we ran PANTHER overrepresentation tests [43] of each using *Anolis carolinensis* as reference with genes annotated in *Laticauda* as a filter.

### Continued expression of *Harbinger-Snek*

To test if *Harbinger-Snek* is expressed in *L. laticaudata* we aligned raw RNA-seq reads from four tissues to the *L. laticaudata* genome from Kishida et al. [21] (BioProject PRJDB7257) using STAR [44] and examined the location of intact *Harbinger-Snek* TEs in IGV [45]and exons in which we had identified *Harbinger* insertions.

## Results and discussion

### *Harbinger-Snek* is unlike transposons seen in terrestrial elapid snakes

Our *ab initio* repeat annotation revealed a novel *Harbinger* DNA transposon in *L. colubrina*, *Harbinger-Snek*. Using BLASTN we found *Harbinger-Snek* present in both *L. colubrina* and *L. laticaudata*, but failed to identify any similar sequences in terrestrial relatives. *Harbingers* are a superfamily of transposons encoding two proteins, a transposase and a Myb-like DNA-binding protein [46]. While both are necessary for transposition [47], we identified multi-copy variants of *Harbinger-Snek* which encoded only one of the two proteins. These variants likely result from large deletions, and may be non-autonomous. In addition, we identified many short non-autonomous variants which retain the same TSDs and terminal motifs, yet encode no proteins.

### *Harbinger-Snek* was horizontally transferred to Laticauda

Harbingers have previously been reported in a wide variety of aquatic vertebrates including fish, crocodilians and testudines, but not in terrestrial vertebrates [29]. Our repeat annotation of the *Laticauda, Aipysurus*, *Naja, Notechis* and *Pseudonaja* assemblies confirmed *Harbinger-Snek* is unique to the two *Laticauda* species examined and is the dominant transposable element in both species (Table 1). This absence from relatives suggested *Harbinger-Snek* was horizontally transferred into the ancestral *Laticauda* genome and our search of over 600 metazoan genome assemblies identified similar sequences only in echinoderms, bivalves and teleosts.

**Table 1:** The expansion of *Harbinger* elements in *Laticauda* spp. This expansion, along with that of LTR elements, in *L. colubrina* has contributed to *L. colubrina having* a larger genome than terrestrial species. This gain in *L. laticaudata* appears to have been offset to some degree by loss from other TE families. Mbp or percentage difference in assembly repeat content between *Laticauda* and the average of the terrestrial *Notechis scutatus* and *Pseudonaja textilis*. Repeat content was annotated using RepeatMasker [39] using a combined Repbase [29] and curated RepeatModeler2 [20] library.

|  | Terrestrial | *L. colubrina* |  | *L. laticaudata* |  |
| --- | --- | --- | --- | --- | --- |
| Retrotransposons |  |  | Diff. Mbp (%) |  | Diff. Mbp (%) |
| SINEs (Mbp) | 25.81 | 24.31 | -1.27 (-0.06%) | 24.57 | -1.00 (-0.06%) |
| Penelopes (Mbp) | 33.19 | 42.34 | +9.20 (0.45%) | 45.28 | +12.15 (0.78%) |
| LINEs (Mbp) | 277.65 | 262.89 | -9.33 (-0.46%) | 235.46 | -36.76 (-2.36%) |
| LTR elements (Mbp) | 175.52 | 202.06 | +27 (1.33%) | 131.33 | -43.73 (-2.81%) |
| DNA transposons |  |  |  |  |  |
| hATs (Mbp) | 88.63 | 79.33 | -6.92 (-0.34%) | 77.62 | -8.63 (-0.55%) |
| Tc1/Mariners (Mbp) | 61.56 | 57.80 | -1.11 (-0.05%) | 55.43 | -3.48 (-0.22%) |
| Harbinger (Mbp) | 0.44 | 229.84 | +229.42 (11.33%) | 126.84 | +126.42 (8.11%) |
| Rolling-circles (Mbp) | 3.24 | 3.09 | -0.13 (-0.01%) | 3.01 | -0.20 (-0.01%) |
| Unclassified (Mbp) | 165.40 | 140.72 | -20.15 (-1.00%) | 134.11 | -26.77 (-1.72%) |
| Total TEs (Mbp) | 798.05 | 999.63 | +217.30 (10.73%) | 788.05 | 5.72 (0.37%) |
| Assembly size (Mbp) | 1,665.53 | 2,024.69 | +396.91 (19.60%) | 1,558.71 | -69.01 (-4.43%) |

The nucleotide sequences most similar to *Harbinger-Snek* were identified in the purple sea urchin, *Strongylocentrotus purpuratus*, and were ~90% identical to the transposase coding region and ~88% identical to the DNA-binding protein. Based on a) high numbers of *Harbinger-Snek* in both species of *Laticauda* sampled and b) similar sequences only present in in marine species, we conclude that *Harbinger-Snek* was likely horizontally transferred to *Laticauda* following their divergence from terrestrial snakes 15-25 Mya, and prior to the crown group divergence of the eight recognised species in *Laticauda* (spanned by *L. colubrina and L. laticaudata*) ~15 Mya [16].

Our phylogenetic analysis (Figure 1) of similar *Harbinger* transposase sequences placed *Harbinger-Snek* in a strongly supported cluster with *Harbingers* found in two sea urchins, *S. purpuratus* and *Hemicentrotus pulcherrimus* (order Echinoida). Interestingly, neither Echinoida assembly contained more than 10 *Harbinger-Snek*-like transposons, none of which encode both proteins. *H. pulcherrimus Harbinger-Snek*-like transposons only contained the transposase, while the *S. purpuratus* assembly contained *Harbinger-Snek*-like transposons encoding either the transposase or the DNA binding protein. In addition, the species that cluster together elsewhere on the tree are not closely related, for example, the sister cluster to the *Laticauda-*Echinoidea cluster contains a variety of fish and bivalve species. The mismatch of the species tree and the transposase tree suggests horizontal transfer of *Harbinger-Snek*-like transposons may be widespread among these marine organisms.

**Figure 1.**
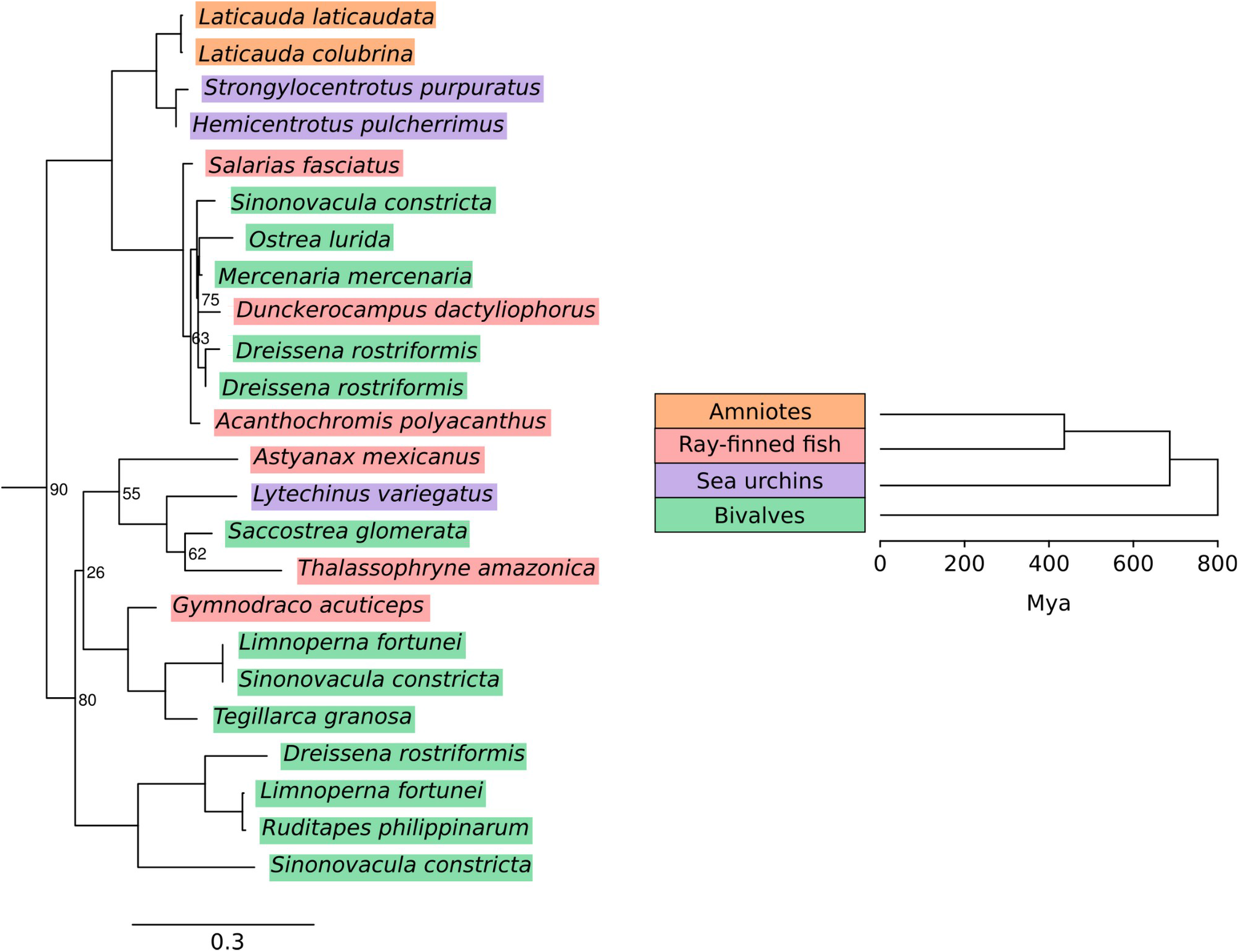
The absence of *Harbinger-Snek* from terrestrial vertebrates and its highest similarity to *Harbingers* present in sea urchins support its horizontal transfer to *Laticauda* since transitioning to a marine habitat. Nodes without support values have support of 95% or higher. The distribution of species across this tree suggests *Harbinger-Snek*-like transposons were horizontally transferred into a wide variety of species. This figure is an extract of a maximum likelihood phylogeny constructed from the aligned nucleotide sequences of the transposases present in curated elements using IQ-TREE 2 [33], for the full tree see SI Figure 1. We also reconstructed trees with similar topologies using RAxML and MrBayes (see methods). Species phylogeny constructed with TimeTree [38].

### *Harbinger-Snek* expanded rapidly in *Laticauda* and is now much less active

Both the RepeatMasker annotation and BLASTN searches reveal a massive expansion in both *Laticauda* species, making up 8% of the *L. laticaudata* assembly and 12% of the larger *L. colubrina* assembly (Table 1, Figure 2). To become established within a host genome following horizontal transfer, TEs must rapidly proliferate, or be lost due to genetic drift or negative selection [48]. To our knowledge the largest previously described expansion of DNA transposons in amniotes following HT is that of *hAT*s in the bat *Myotis lucifugus* [13–15]. Following HT ~30 Mya, *hAT* transposons quickly expanded over 15 My at an estimated rate of ~0.7 Mbp/My and currently make up ~3.3% of the *M. lucifugus* genome. Using the upper bound of *Harbinger-Snek*’s transfer of 25 My (directly after their divergence from terrestrial Australian snakes), we calculate *Harbinger-Snek* to have expanded in *L. colubrina* at a rate of 11.3 Mbp/My and in *L. laticauda* a rate of 8.12 Mby/My. Therefore, our finding is the largest described expansion of a TE in an amniote following HTT.

**Figure 2.**
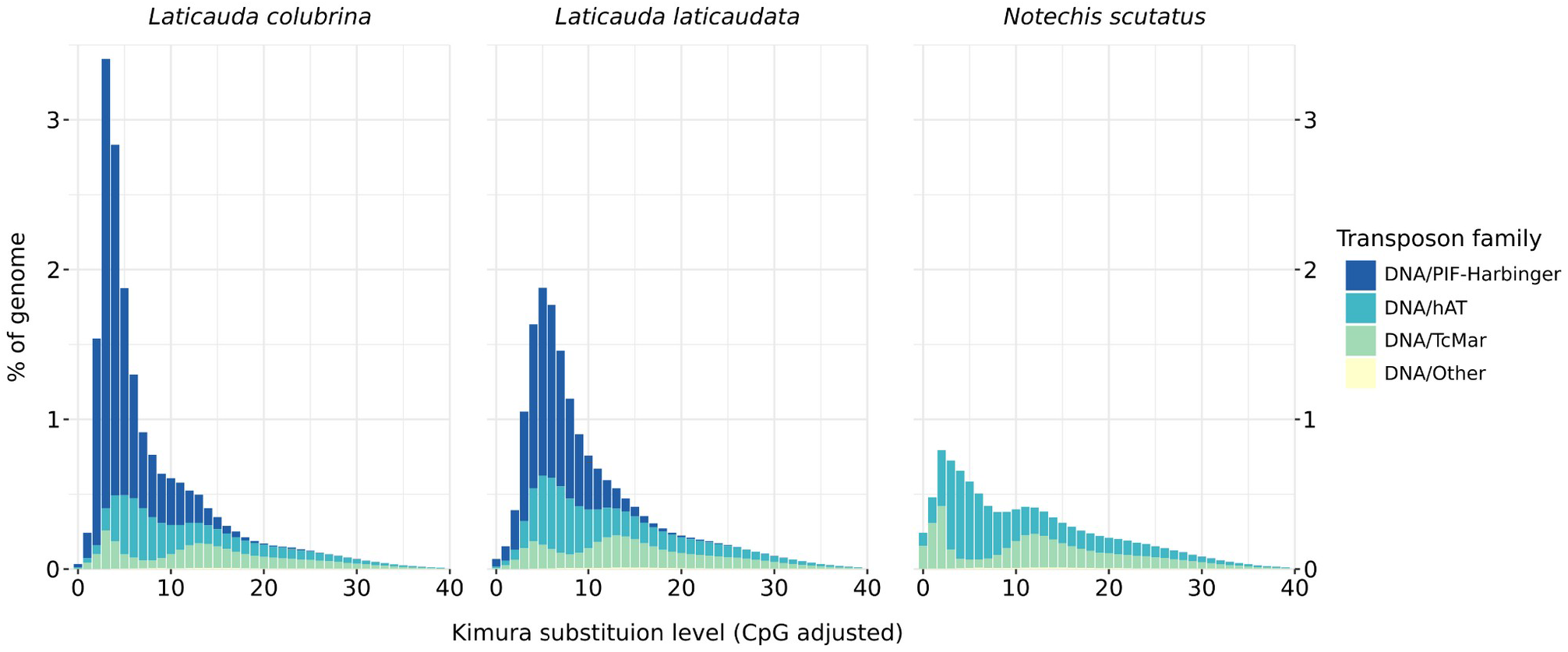
Rapid, recent expansion of *Harbinger-Snek* PIF-Harbinger transposons. Horizontal transfer of this transposon into the *Laticauda* ancestor has occurred within the past 15-25 My [16]. Due to expansions since then, these transposons have become the dominant DNA transposon in *Laticauda* genomes, in contrast to the genomes of their closest terrestrial relatives such as *Notechis scutatus* (diverged ~15-25 Mya). Repeat content calculated with RepeatMasker [39].

Mass expansion of existing TEs during speciation has previously been seen in many groups including primates [49], woodpeckers [50] and salmonids [51]. By making the genome more dynamic these expansions fostered rapid adaptations. The sharp peak in the divergence profile (Figure 2) indicates *Harbinger-Snek’s* expansion was rapid, and the small number of near-identical copies suggests expansion has slowed massively, especially in *L. laticaudata*. Many more copies of *Harbinger-Snek* able to transpose are present in the *L. colubrina* assembly than the *L. laticaudata* assembly, with only 1 fully intact copy in *L. laticaudata*, but 269 in *L. colubrina*. Our alignment of *L. laticaudata* RNA-seq data from four tissues (vomeronasal organ, nasal cavity, tongue and liver) to the *L. laticaudata* genome revealed reads mapping across both coding regions of the intact copy of *Harbinger-Snek*. Therefore, *Harbinger-Snek* and its non-autonomous derivatives may still be transposing in *L. laticaudata*.

In addition to containing many more intact copies of the full element, *Laticauda colubrina* also contains a higher number of the aforementioned “solo-ORF” variants than *L. laticaudata*, with 2263 intact transposase only variants compared to 35, and 452 intact DNA binding protein only variants compared to 6. Based on this stark contrast, since divergence ~15 Mya [16] either *L. colubrina* has maintained a higher rate of *Harbinger-Snek* expansion or *L. laticaudata* has had a higher rate of *Harbinger-Snek* loss; or more likely, a combination of these two effects.

### The accordion model - the expansion of *Harbinger-Snek* has been balanced by loss in *L. laticaudata*

The peak in *Harbinger-Snek* expansion in *L. colubrina* is both higher and more recent than *L. laticaudata* (Figure 2). In addition *L. laticaudata* has a much lower overall *Harbinger-Snek* content and genome size (Table 1). Past observations in birds, mammals and squamates found increases in genome size due to transposon expansion are balanced by loss due to deletions through non-allelic homologous recombination (NAHR) [52,53]. We expect that the mass expansion of *Harbinger-Snek* in *Laticauda* has generated many near identical sites in the genome, in turn promoting NAHR. In spite of the explosive expansion of *Harbinger-Snek* in *L. laticaudata*, the genome size and total TE content is very similar to that of the terrestrial *Pseudonaja* and *Notechis* (Table 1). This retention of a similar genome size is not seen in *L. colubrina*, the genome assembly of which is 20% larger than the terrestrial species. However, the overall TE content of the *L. colubrina* genome remains similar to that of *L. laticaudata* and the terrestrial species, with the expansion of TEs only contributing half of the total increase in genome size. This is consistent with the aforementioned balancing of TE expansion by deletions.

### Expansion of *Harbinger-Snek* has potentially impacted gene function

In both species of *Laticauda* many insertions of *Harbinger-Snek* overlap with or are contained within exons, regulatory regions and introns. Insertions overlapped with the exons of 56 genes in *L. colubrina* and 31 in *L. laticaudata*, 17 of which are shared (SI Table 1). By manually inspecting transcripts mapped to the *L. laticaudata* genome we determined 8 3’ UTRs and 2 coding exons predicted by Liftoff now contain *Harbinger-Snek* insertions which contribute to mRNA (SI Table 1). These genes have a wide range of functions, many of which could be significant in the context of adaptation. We also identified insertions into 1685 and 888 potentially regulatory regions (within 5 kbp of the 5’ UTR in genes) and into introns of 4141 and 1440 genes in *L. colubrina* and *L. laticauda* respectively. PANTHER over/under-representation tests of these in gene and regulatory region insertions identified a number of pathways of potential adaptive significance (SI Tables 2-5). Therefore, *Harbinger-Snek* is a prime candidate in the search for genomic changes responsible for *Laticauda’s* adaptation to a marine environment through altered gene expression.

## Conclusion

In this report, we describe the rapid expansions of *Harbinger-Snek* TEs in *Laticauda* spp., to our knowledge, the fastest expansion of a DNA transposon in amniotes reported to date. The large number of insertions of *Harbinger-Snek* into exons and regulatory regions may have contributed to speciation and adaptation to a new habitat; this suggests a number of future lines of investigation. As the HTT was prior to the divergence of 8 *Laticauda* species, *Harbinger-Snek* presents a unique opportunity to reconstruct subsequent molecular evolution and determine the impact of HTT on the adaptation of *Laticauda* to the amphibious-marine habitat.

## Supplementary Information

SI Table 1 - *Laticauda colubrina and Laticauda laticaudata* genes with *Harbinger-Snek* insertions into or overlapping open reading frames, and any noticeable effects on insertion noted from transcript data. Gene coordinates predicted with Liftoff [40] using the RefSeq *Notechis scutatus* assembly and gene annotation as reference. Repeat annotation performed with RepeatMasker [39] using a custom repeat library (see Methods). Intersect performed using BEDTools [54]. Transcripts mapped to the genome assembly using STAR [44] and viewed in IGV [45].

SI Table 2 - Biological processes with an over/under-representation of *Harbinger-Snek* insertions into *Laticauda colubrina* genes. Representation test performed using PANTHER [43]. Gene coordinates predicted with Liftoff [40] using the RefSeq *Notechis scutatus* assembly and gene annotation as reference. Repeat annotation performed with RepeatMasker [39] using a custom repeat library (see Methods). Intersect performed using plyranges [42].

SI Table 3 - Molecular functions with an over/under-representation of *Harbinger-Snek* insertions into *Laticauda colubrina* genes. Representation test performed using PANTHER [43]. Gene coordinates predicted with Liftoff [40] using the RefSeq *Notechis scutatus* assembly and gene annotation as reference. Repeat annotation performed with RepeatMasker [39] using a custom repeat library (see Methods). Intersect performed using plyranges [42].

SI Table 4 - Biological processes with an over/under-representation of *Harbinger-Snek* insertions into potential regulatory regions of *Laticauda colubrina* genes. Representation test performed using PANTHER [43]. Gene coordinates predicted with Liftoff [40] using the RefSeq *Notechis scutatus* assembly and gene annotation as reference. Repeat annotation performed with RepeatMasker [39] using a custom repeat library (see Methods). Intersect performed using plyranges [42].

SI Table 5 - Molecular functions with an over/under-representation of *Harbinger-Snek* insertions into potential regulatory regions of *Laticauda colubrina* genes. Representation test performed using PANTHER [43]. Gene coordinates predicted with Liftoff [40] using the RefSeq *Notechis scutatus* assembly and gene annotation as reference. Repeat annotation performed with RepeatMasker [39] using a custom repeat library (see Methods). Intersect performed using plyranges [42].

SI Table 6 - Latin species names and versions of all public genomes used. All were downloaded from RefSeq [41] when available, else from GenBank [55].

